# Criticality of resting-state EEG predicts perturbational complexity and level of consciousness during anesthesia

**DOI:** 10.1101/2023.10.26.564247

**Authors:** Charlotte Maschke, Jordan O’Byrne, Michele Angelo Colombo, Melanie Boly, Olivia Gosseries, Steven Laureys, Mario Rosanova, Karim Jerbi, Stefanie Blain-Moraes

## Abstract

1

Consciousness has been proposed to be supported by electrophysiological patterns poised at criticality, a dynamical regime which exhibits adaptive computational properties, maximally complex patterns and divergent sensitivity to perturbation. Here, we investigated dynamical properties of the resting-state electroencephalogram of healthy subjects undergoing general anesthesia with propofol, xenon or ketamine. We then studied the relation of these dynamic properties with the perturbational complexity index (PCI), which has shown remarkably high sensitivity in detecting consciousness independent of behavior. All participants were unresponsive under anesthesia, while consciousness was retained only during ketamine anesthesia (in the form of vivid dreams)., enabling an experimental dissociation between unresponsiveness and unconsciousness. We estimated (i) avalanche criticality, (ii) chaoticity, and (iii) criticality-related measures, and found that states of unconsciousness were characterized by a distancing from both the edge of activity propagation and the edge of chaos. We were then able to predict individual subjects’ PCI (i.e., PCI_max_) with a mean absolute error below 7%. Our results establish a firm link between the PCI and criticality and provide further evidence for the role of criticality in the emergence of consciousness.

**Significance Statement:** Complexity has long been of interest in consciousness science and had a fundamental impact on many of today’s theories of consciousness. The perturbational complexity index (PCI) uses the complexity of the brain’s response to cortical perturbations to quantify the presence of consciousness. We propose criticality as a unifying framework underlying maximal complexity and sensitivity to perturbation in the conscious brain. We demonstrate that criticality measures derived from resting-state electroencephalography can distinguish conscious from unconscious states, using propofol, xenon and ketamine anesthesia, and from these measures we were able to predict the PCI with a mean error below 7%. Our results support the hypothesis that critical brain dynamics are implicated in the emergence of consciousness and may provide new directions for the assessment of consciousness.

## Introduction

Neuroscience is increasingly borrowing from complex systems theory in order to understand the link between neural dynamics, behavior and conscious states. In nature, complexity often emerges in systems poised between two dynamical regimes such as chaos and stability—a phenomenon known as criticality (1, 2). At this fine balance point, systems display optimal computational capacity, maximally complex patterns, and divergent sensitivity to external perturbation. In virtue of these features, criticality is increasingly explored as a requirement for healthy brain function (2–4) and the emergence of consciousness (5–8).

Although the association of criticality with consciousness is rather recent, complexity has long been of interest in consciousness science. Early theoretical work suggested that consciousness is tightly linked to “neural complexity”, which measures the balance between functional differentiation and integration within a system (9), an idea that gave rise to the Integrated Information Theory (IIT) of consciousness (10, 11). A variety of other complexity measures, based in various theoretical paradigms, have been identified as markers of consciousness in physiological, pharmacological and pathological conditions (7, 12–17).

Among these measures, the perturbational complexity index (PCI) (18) captures the complexity of the brain’s response to a direct and non-invasive cortical perturbation using transcranial magnetic stimulation (TMS) and electroencephalography (EEG). Due to its unique sensitivity in detecting consciousness in patients affected by disorders of consciousness, it stands today a promising index for the assessment of human consciousness (15, 19, 20). The question then arises as to which properties of the conscious brain underpin high PCI. Knowledge of these properties may not only inform theories of consciousness but may also point the way to new clinical measures of consciousness that do not require a TMS machine – a device with only scant accessibility in clinical. Originally, PCI was inspired by IIT and the concept of integration-differentiation balance. However, the link between PCI and IIT is not exclusive (21, 22), allowing for alternative or complementary explanatory theories.

A natural explanation may be found in criticality. The complexity of evoked responses, as measured in PCI, is in fact predicted to be maximal in systems poised at criticality (23–25). As such, criticality has been proposed as a unifying framework underlying maximal complexity and sensitivity to perturbation in the conscious brain (2, 5, 7, 26). Still, while previous studies have suggested a conceptual link between criticality and maximally integrated information (6, 27), the relation between the PCI and criticality of the pre-TMS resting-state EEG remains unexplored. In this study, we investigate whether criticality measures derived from resting-state EEG (without TMS) can distinguish conscious from unconscious states in a pharmacological model of (un)consciousness using propofol, xenon and ketamine anesthesia. Moreover, we explored the potential of these measures to predict the PCI value (i.e., PCI_max_), aiming at shedding light on the physical bases of this index.

Brain criticality has been approached through a diverse set of perspectives and methods (2 as a review, 6, 7, 28, 29). Here, we explore measures of two types of criticality: 1) avalanche criticality and 2) the edge of chaos (see Methods, see Fig.7). Both types of criticality describe the meeting point of two dynamical regimes, namely 1) activity amplification and dissipation and 2) chaos and stability, respectively. In addition, we analyzed a set of ‘criticality-related measures’ - a group of properties that are associated with criticality in general but that are not known to be a specific feature of any one criticality type (e.g., Lempel-Ziv complexity).

In the first part of this study, we describe the effect on brain criticality of general anesthesia with propofol, xenon or ketamine. Each anesthetic procedure was tailored to reach a common behavioral state of unresponsiveness—in other words, a ‘surgical level’ of general anesthesia, delivered to healthy participants in the absence of any surgery. While all participants were behaviorally unresponsive during drug exposure, only anesthesia with ketamine led to a clear-cut dissociation between responsiveness and consciousness (30, 31), with subjects being unresponsive while also having intense conscious experience (also known as ketamine dreams) (see 15 for example reports). In the second part, we examine the relation between resting-state brain criticality just prior to TMS perturbation and the complexity of the response immediately following the TMS perturbation (i.e., PCI).

We hypothesized that states of unconsciousness (i.e., during general anesthesia with propofol or xenon) diverge from criticality, either to the sub- or supercritical state, and that the brain exhibits close-to-critical dynamics only when consciousness is present (see Fig.1). Meanwhile, general anesthesia with ketamine was not expected to induce a deviation from criticality, but rather to maintain close-to-critical dynamics, similarly to normal wakefulness. We further hypothesized that the level of criticality of resting-state cortical activity can predict the complexity of the response to perturbation using TMS (i.e., PCI). Whereas brains poised at criticality were expected to show a highly complex reaction to the targeted perturbation (i.e., high PCI), sub- and supercritical dynamics were expected to display a local and quickly vanishing reaction (i.e., low PCI), or a wide-ranging but stereotypical reaction (also low PCI), respectively (see Fig.1). Our objective is to provide a mechanistic framework for one of today’s most reliable metrics of consciousness, PCI, from which we may derive a new and complementary approach for the assessment of consciousness.

**Figure 1:**
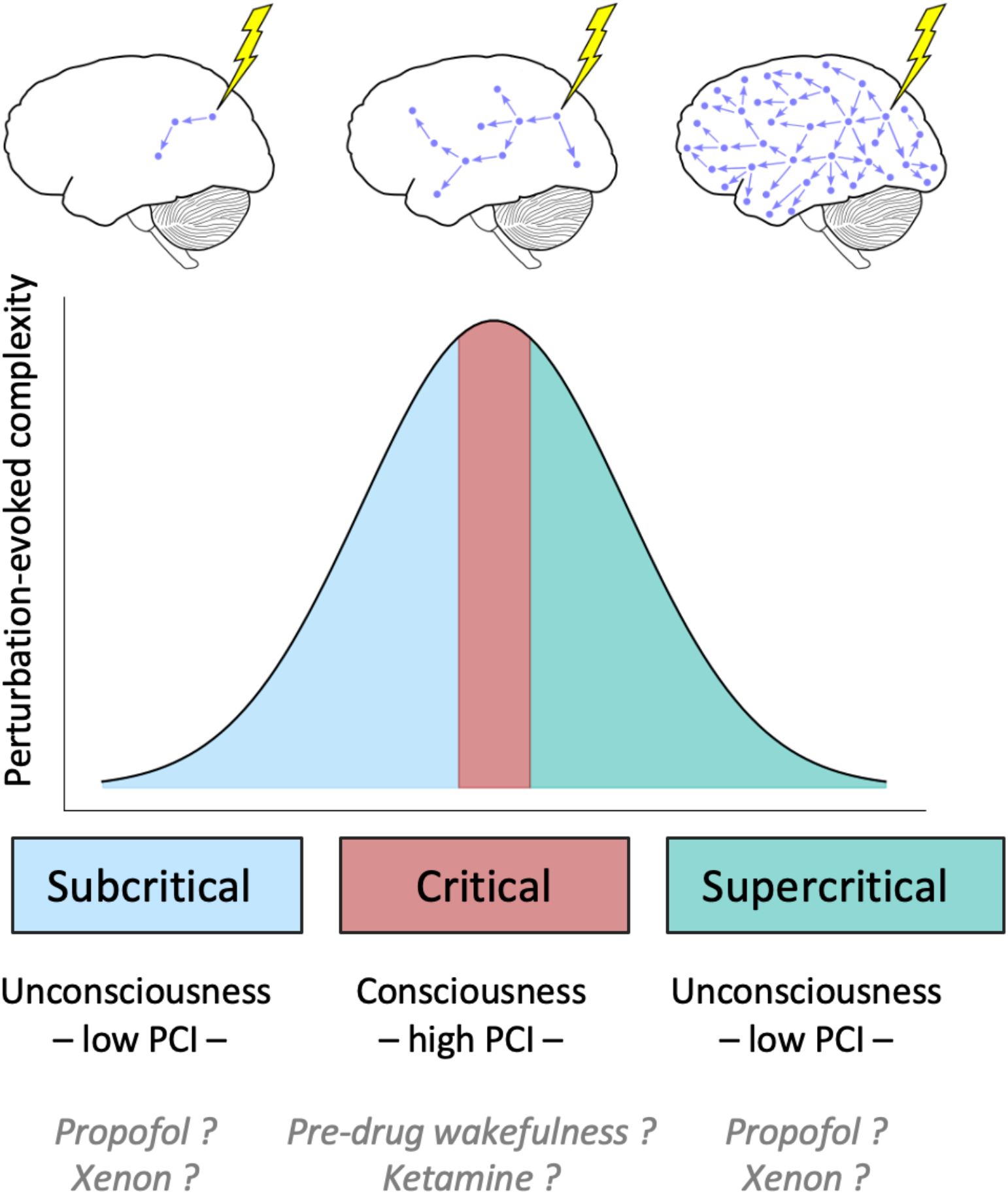
Illustration of hypothesis: A system at criticality is poised between two dynamical regimes and exhibits adaptive computational properties, including maximally complex patterns and divergent sensitivity to perturbations. As such, criticality offers a suitable framework for explaining the perturbation-evoked complexity measured by PCI. The top row of the figure illustrates the concept of avalanche criticality. Arrows indicate activity propagation over time resulting from a single perturbation (e.g., a sensory event or a somatic signal). **(Top Left)** In a subcritical regime, a single unit activation or event triggers on average less than one additional event (branching ratio < 1). Thus, the effect of a single perturbation quickly dissipates and has no long-term (time) or long-range (space) effect on the system. In other words, the system is highly stable and quickly ‘forgets’ information about its inputs. **(Top Right)** In a supercritical regime, a single event triggers on average more than one downstream event (branching ratio > 1). The effect of a single perturbation exponentiates quickly over time leading to total activation of the system. The system is thus highly unstable, and the over-amplification of signals results in rapid forgetting through information corruption. **(Top Middle)** At criticality, a single event triggers exactly one downstream event on average (branching ratio = 1). Variations around this average yield a diverse set of network responses of all sizes and durations, facilitating communication between the system’s microscopic and macroscropic scales. The system is poised between stability and instability (balancing reliability and flexibility), and information reverberates across the system and over prolonged timespans. **(Bottom)** We hypothesize that states of consciousness (i.e., normal wakefulness and ketamine anesthesia) are poised at criticality. States of unconsciousness (i.e., during general anesthesia with propofol or xenon) are hypothesized to diverge from criticality, either to a sub- or supercritical state. We further hypothesize that sensitivity to perturbations (i.e., complexity of the response to external stimulation), as quantified by the perturbational complexity index (PCI) is maximized at criticality and reduced in sub- and supercritical states.

## 3 Results

We analyzed data from a previously published study (15, 32), consisting of 15 healthy adults who were exposed to propofol (*n*=5), xenon (*n*=5) or ketamine (*n*=5) general anesthesia. Spontaneous electroencephalography (EEG) was recorded (~5 min) during resting wakefulness prior to drug administration and during drug-induced loss of responsiveness (see 15 for full protocol). The PCI values (PCI_max_) for every subject before and during drug administration were obtained by Sarasso et al. (15) using a TMS-EEG protocol (18). Although drug administration resulted in unresponsiveness in all three groups, only participants exposed to propofol or xenon were considered unconscious (e.g., did not report any subjective experience). In the ketamine condition, participants reported vivid, conscious dream-like experiences upon recovery of responsiveness (15).

### 3.1 Propofol and xenon, but not ketamine, induce a shift away from avalanche criticality

For a large class of dynamical systems, activity spreads through so-called *avalanches —* “chain reactions” or cascades of activity that propagate through time and space. In ordered, subcritical systems, avalanches tend to be short-lived with a characteristic small scale, whereas in disordered, supercritical systems, a large number of avalanches span the whole system, again imposing a characteristic (system-size) scale to the avalanche distribution. In contrast, at the avalanche-critical point, avalanches are scale-free — no scale dominates, such that the probability distribution of their features, such as size and duration, converge on a power law (the only scale-invariant mathematical function). Therefore, the presence of power law distributions of avalanche statistics constitutes a first indicator of avalanche-critical dynamics.

Avalanche detection on EEG data requires binarization of the signal, using a threshold of *n* standard deviations (SD). Following other studies (33, 34), the optimal threshold for avalanche detection was identified by finding the point of divergence between the probability distribution of z-scored EEG signal values and a best-fit Gaussian (Fig. 2B). For comparison, the corresponding probability distribution for absolute (non-z-scored) EEG signal values is shown in Fig. 2A. Note that although the amplitude excursions are wider for xenon and propofol conditions in terms of raw microvolt values (Fig. 2A), z-scoring reveals that the shape of the distribution is substantially more heavy-tailed for the wakefulness and ketamine conditions (Fig. 2B), consistent with previous results from invasive electrocorticography recordings in nonhuman primates (34). Such heavy-tailed distributions are a hallmark of critical dynamics. The point of divergence between the Gaussian and the observed data was estimated at 2.0 SD (see Fig 2C) and was taken as a threshold for detecting non-stochastic neural events (i.e., for binarization). From the binarized signal, avalanches were detected using an inter-event interval of 8 ms. All results were replicated on a range of hyperparameters (1.5-3.0 SD, 4-12ms) and are provided in Supplementary Material 1.

**Figure 2:**
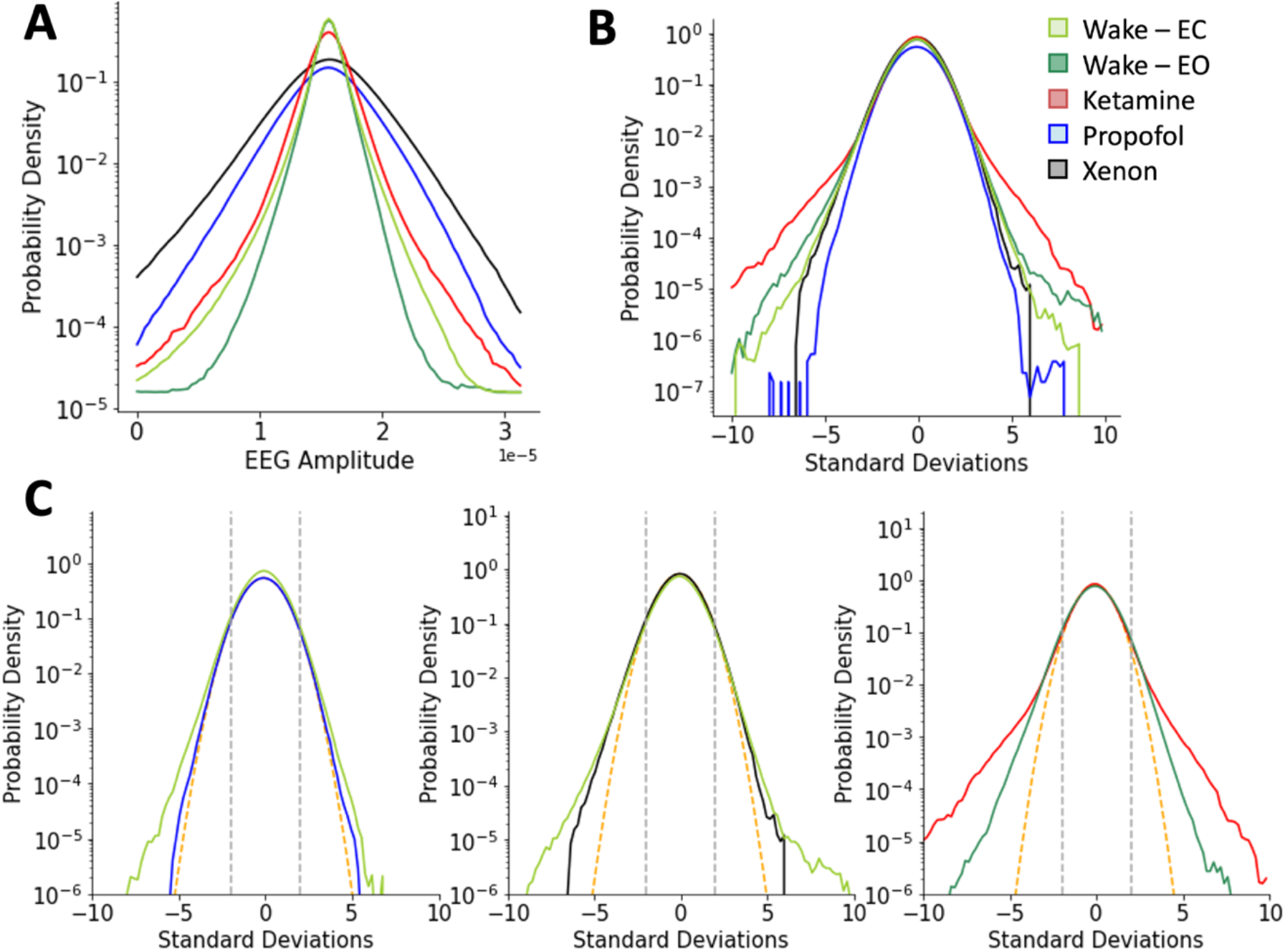
Probability distribution of signal values in terms of amplitude and standard deviation. Distributions were estimated over all channels and averaged across participants. The point of divergence between these distributions and their best-fit Gaussian marks a boundary beyond which observed fluctuations are unlikely to be the result of stochastic variability around the mean and can thus be considered as ‘neural events’. **(A)** Probability distribution of amplitudes by condition. Signal excursions are broadest in the propofol (blue) and xenon (black) conditions and narrower during pre-drug wakefulness (light green, eyes closed; dark green, eyes open), and during ketamine anesthesia (red) **(B)** Probability distribution of z-scores by condition. While propofol (blue) and xenon (black) distributions vanish faster, ketamine (red) and wakefulness (light green, eyes closed; dark green, eyes open) exhibit a heavy-tailed distribution, suggestive of avalanche dynamics. For visualization only, the wakefulness eyes-open condition was plotted as the average over all 10 subjects. **(C)** Whereas propofol (blue) and xenon (black) more closely follow a Gaussian distribution, ketamine (red) and wakefulness (light green, eyes closed; dark green, eyes open) deviate from the Gaussian (orange dashed curve) above the observed threshold of 2 SD (grey dashed line). Each line corresponds to the average over 5 subjects in the given condition.

Despite larger absolute amplitudes during propofol (*t*(4) = -5.71, *p*<0.01) and xenon (*t*(4) = -5.04, *p*<0.01) anesthesia (see Fig. 2A), both conditions exhibited significantly fewer avalanches compared to wakefulness (propofol *t*(4) = -4.00, *p*<0.05, xenon: *t*(4) = -4.06, *p*<0.05). In contrast, the number of avalanches remained unchanged during ketamine anesthesia. The distribution of avalanche sizes followed a truncated power law with exponential drop-off in 25 out of 30 recordings (10/15 during wakefulness, 15/15 during anesthesia recordings) (see Supplementary Methods, Note 3), while the distribution of avalanche durations followed a truncated power law with exponential drop-off in all recordings across participants and conditions. An inspection of the avalanche distribution (i.e., visualized using the complementary cumulative distribution function) (Fig 3A) reveals that exposure to xenon and propofol yields an earlier exponential drop-off, suggestive of subcritical dynamics. Most interestingly, ketamine showed a distribution more similar to wakefulness, which suggests dynamics closer to criticality than xenon and propofol, but less critical than wakefulness. These differences are especially clear in the avalanche size distributions (Fig 3 A, left panel).

**Figure 3:**
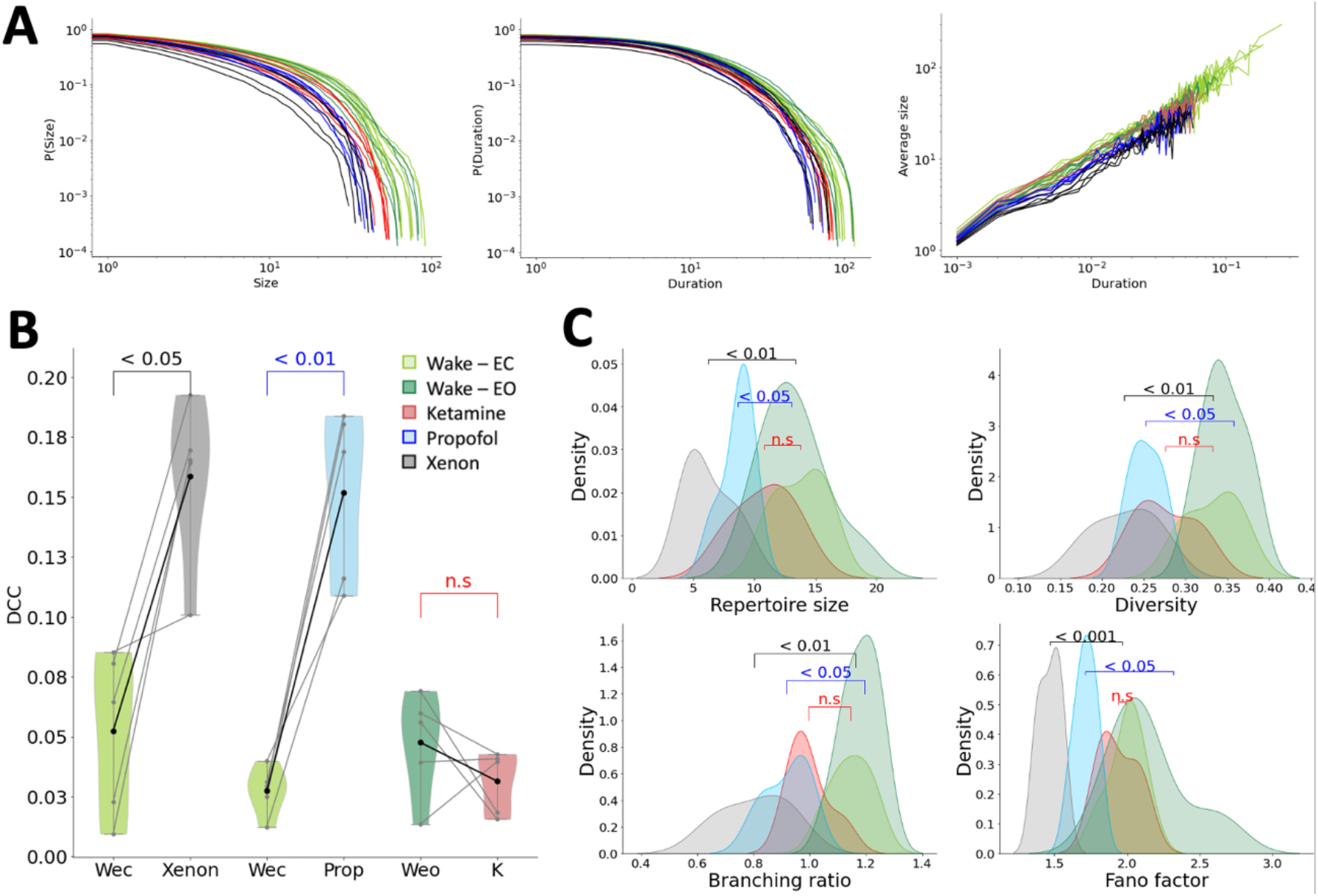
**A)** Distribution of avalanche size (left), duration (middle) and average size for a given duration (right), visualized using the complementary cumulative distribution function for each subject and condition individually: xenon (black), propofol (blue), ketamine (red), wakefulness with eyes open (‘Wake-Eo’, dark green) and closed (‘Wake-Ec’, light green). In all three distributions, the propofol and xenon conditions exhibit an earlier drop-off, compared to wakefulness. The ketamine condition is intermediate between wakefulness and unresponsive anesthesia (xenon and propofol) conditions, indicating that large and sustained avalanches are more strongly maintained during exposure to ketamine than they are under xenon or propofol. **B)** The deviation from criticality coefficient (DCC) increases under the effect of xenon and propofol, but not ketamine. Light grey lines represent individual subjects, bold lines are the mean over subjects. **C)** Effect of xenon, propofol and ketamine on measures of criticality: xenon and propofol but not ketamine resulted in decreases in the size and diversity of the avalanche pattern repertoire, the branching ratio and the Fano factor, each of which is expected for more subcritical dynamics.

These effects can be approximately quantified by comparing the slopes of the best-fit power laws, as the steepness of the best-fit line is expected to increase as a monotonic function of the degree of subcriticality (see Supplementary Note 2). We therefore used the best-fit power-law slope as an indirect metric of the distance from criticality. We further quantified the likelihood that the distributions followed a power law by comparing goodness of fit between power law, lognormal and exponential functions using loglikelihood estimation (36, 37).

Compared to wakefulness, the administration of propofol (t(4) = -2.88, p<0.05), xenon (t(4) = -3.04, p<0.05) and ketamine (t(4) = -4.23, p<0.05) significantly increased the slope of the best-fit power law of avalanche duration, indicating the occurrence of overall shorter-lived avalanches during drug exposure (see Fig 3A). The slope for avalanche size also increased during exposure to propofol (t(4) = -7.18, p<0.01) and xenon (t(4) = -4.93, p<0.05) but not ketamine, indicating a decrease of large-sized avalanches during unconsciousness (see Fig 3A). The slope of the distribution of average size by duration increased upon administration of propofol (t(4) = -4.59, p<0.05), xenon (t(4) = -6.68, p<0.01) and ketamine (t(4) = -4.44, p<0.05). For the distribution of avalanche sizes, the likelihood of a power law behavior significantly decreased during general anesthesia with propofol (t(4) = 4.91, p<0.05) and xenon (t(4) = 12.13, p<0.01), but not ketamine. For the distribution of avalanche duration, only exposure to xenon (t(4) = 5.22, p<0.01) significantly decreased the likelihood of a power law behavior. The decrease in propofol did not reach significance.

Taken together, exposure to propofol, xenon and ketamine yielded a decreased probability of large and long-lasting avalanches, and a deviation from power-law behavior, providing evidence for subcritical dynamics. While this effect is strongly expressed during propofol- and xenon-induced unconsciousness, exposure to ketamine yielded overall smaller deviations from wakefulness (see Fig. 2A).

While the change in exponents provides preliminary evidence for alterations in the underlying system’s dynamics, a critical system should exhibit a specific relation between its power-law exponents (slopes), which are known as *critical exponents* (35) (29). The observed error of this *scaling relation* is expressed in the deviation from criticality coefficient (DCC, see Methods) (38). Whereas the administration of propofol (t(4) = -7.45, p<0.01) and xenon (t(4) = -4.33, p<0.05) resulted in a large DCC, ketamine did not significantly alter DCC with respect to wakefulness (see Fig 3 B). This supports our hypothesis that only exposure to propofol or xenon shifts neuronal dynamics away from criticality, while exposure to ketamine yields near-critical dynamics that are indistinguishable from wakefulness.

A similar behavior was clearly observed in a variety of other measures of avalanche criticality, namely the branching ratio, the Fano factor and the size and average diversity of the avalanche pattern repertoire (see Fig 3 C). Briefly, the branching ratio estimates the number of events in the next time bin that are expected to arise from a single event in the present time bin, and should be near 1.0 at criticality and smaller for subcritical systems (28). The Fano factor is a measure of the magnitude of fluctuation of the activity signal and is expected to exceed 1.0 at criticality (39). The avalanche pattern repertoire is the set of unique spatial patterns spanned by the observed avalanches (40), and is expected to be maximal in size and diversity at criticality. Each of these measures showed signs of a shift towards subcriticality in the xenon and propofol conditions, but not under ketamine. Specifically, the branching ratio, Fano factor and average repertoire size and diversity all decreased under propofol (branching ratio: *t*(4) = 5.26, *p*<0.05; Fano factor: *t*(4) = 6.18, *p*<0.05; repertoire size: *t*(4) = 3.17, *p*<0.05; repertoire diversity: *t*(4) = 5.22, *p*<0.05) and xenon (branching ratio: *t*(4) = 10.73, *p*<0.01; Fano factor: *t*(4) = 20.50, *p*<0.001; repertoire size: *t*(4) = 6.35, *p*<0.01; repertoire diversity: *t*(4) = 8.86, *p*<0.01).

In summary, unconsciousness following exposure to propofol or xenon yielded network dynamics diverging from criticality into the subcritical phase. Specifically, drug-induced unconsciousness, despite overall larger amplitudes, was characterized by more dissipative activity propagation, smaller activity fluctuations (i.e., heavier-tailed signal distributions) and less diverse avalanches. In contrast, critical dynamics and related network properties (i.e., stable activity propagation, large fluctuations and diverse avalanches) that were observed during wakefulness were preserved during exposure to ketamine. Cumulatively, this evidence strongly suggests that propofol and xenon, but not ketamine, shift neuronal dynamics away from avalanche criticality.

### 3.2 Propofol and xenon, but not ketamine, increase brain chaoticity

Chaos is broadly defined as the sensitivity of a system’s trajectories in phase space to the details of its initial conditions. The edge of chaos marks the turning point where a system switches from dynamics that converge onto fixed-point or periodic attractors to dynamics that wander off into chaos. The edge of chaos exists as its own critical phase transition -- dissociable from avalanche criticality (41, 42) though sharing many high-level properties with it, including maximal signal diversity and sensitivity to perturbation (2). The degree of chaos, or chaoticity, was estimated using three measures: 1) the modified 0-1 chaos test (7, 43) 2) the largest Lyapunov exponent (LLE) (44) and; 3) the standard deviation of the integrated time-lagged covariance matrix, as proposed by Dahmen et al. (41) (see Methods). The 0-1 chaos test and the LLE each were estimated on non-overlapping 10 s windows of signal on each channel individually and averaged over time. The 0-1 chaos test was previously validated on electrophysiological signal that was low-pass filtered at the lowest oscillatory peak (up to 6 Hz), channel-wise (7). However, ketamine anesthesia shows an absence of low-frequency oscillations (15). Thus, we instead applied a fixed low-pass frequency filter across all recordings and repeated the analysis for a range of low-pass frequencies (1-12 Hz with a 1-Hz step; Fig 4 A, bottom).

**Figure 4.**
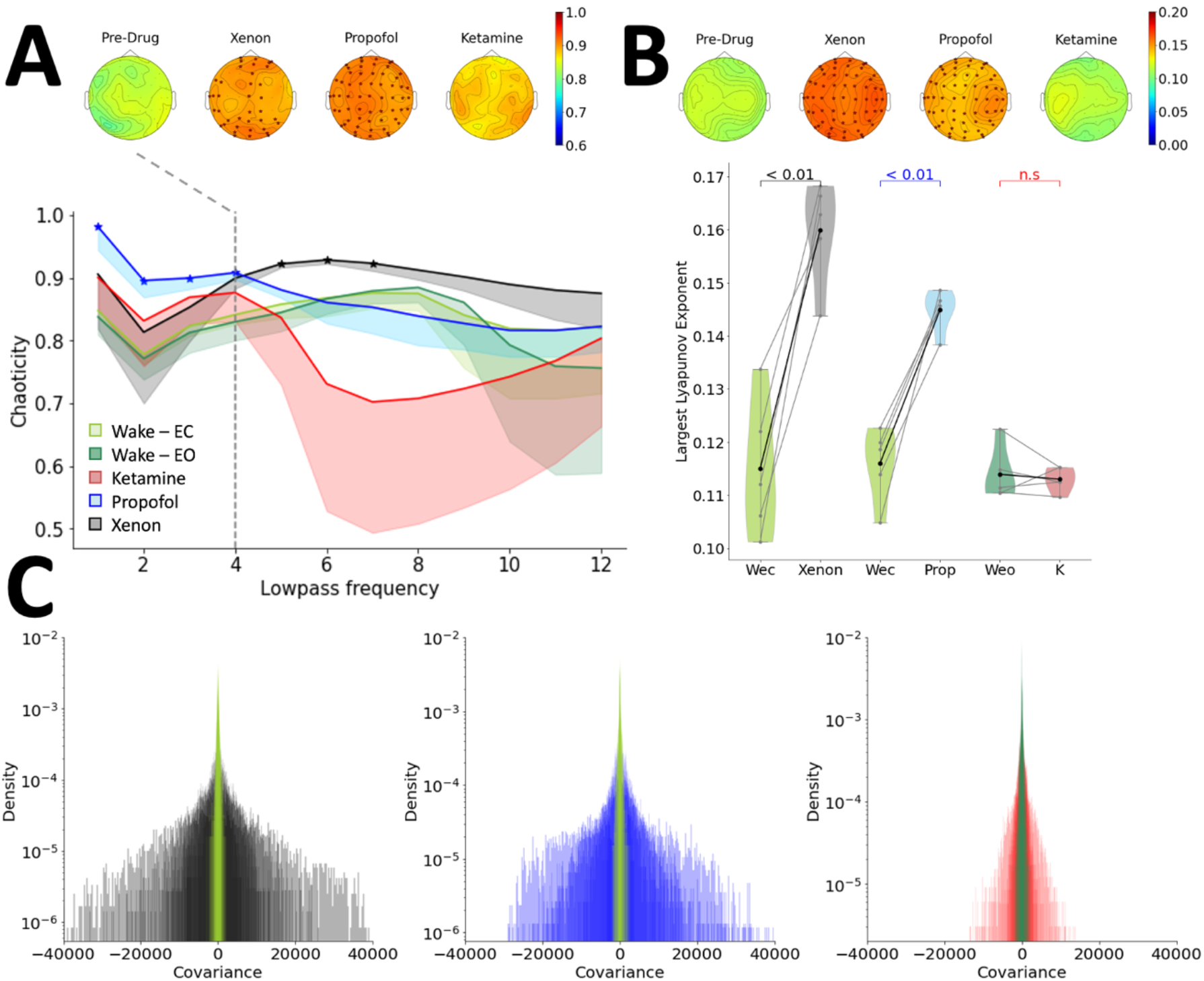
Effects of propofol, xenon and ketamine on brain chaoticity. **A)** Bottom: Chaoticity estimated over a range of low-pass filters, using the 0-1 chaos test. Stars indicate significant differences with the corresponding wakefulness data. Bold lines represent the chaoticity values over all channels, averaged over 5 participants in one condition. The shaded area below the line represents the lower standard deviation. Propofol (blue) significantly increased chaoticity for frequencies up to 4 Hz. Xenon (black) showed a significant increase at low-pass frequencies between 4 and 7 Hz. Top: Topographic distribution visualized for channel-wise chaoticity values (0-1 chaos) with a low-pass filter of 4 Hz. Channels with a significant increase in chaoticity compared to the corresponding wakefulness data are marked with a red star. **B)** Bottom: The largest Lyapunov exponent (LLE) increased during exposure to propofol (blue) and xenon (black), but not ketamine (red). Light grey lines represent individual subjects and bold lines are the mean over subjects. Top: Topographic maps represent the time-averaged LLE for every channel. Channels with a significant increase in chaoticity compared to the corresponding wakefulness data (green) are marked with a red star. **C)** Distribution of integrated time-lagged covariances during propofol (blue), xenon (black) and ketamine (red) anesthetic conditions, compared to the corresponding wakefulness data (green).

Chaoticity of low-frequency dynamics increased following exposure to propofol and xenon. Whereas the propofol-induced increase of chaoticity was most present when using low-pass filters of 1 to 4 Hz (all *P <* 0.05), the xenon-induced increase only occurred when including higher frequencies up to 7 Hz (i.e., using low-pass filters of 4 to 7 Hz) (all *P <* 0.05). Only at a low-pass frequency of 4 Hz was there an increase in chaoticity observed for both drugs. While a propofol-induced increase in chaoticity was observed homogenously over all areas, xenon mostly increased frontal and occipital chaoticity (see Fig 4 A, top). Importantly, no difference in chaoticity was observed between wakefulness and exposure to ketamine. For direct comparison with previous work, we also applied 0-1 chaos using the above-described peak detection method (see Supplementary Material 2).

The LLE was estimated on the broadband signal (1-40 Hz). For comparison between conditions, LLE values were averaged across channels to yield one average LLE per participant and condition. In accordance with the 0-1 chaos findings, the LLE increased during exposure to propofol (*t*(4) = -7.56, *p*<0.01) and xenon (*t*(4) = -6.65, *p*<0.01), but not ketamine. Furthermore, observed increases in chaoticity occurred homogenously over all channels (see Fig 4 B). Similarly, propofol (*t*(4) = -2.96, *p*<0.05) and xenon (*t*(4) = -2.81, *p*<0.05), but not ketamine, significantly increased the width of the covariance matrix (see Fig 4 B), which is indicative of increased chaoticity according to statistical physical models (41).

Altogether, the three measures of brain chaoticity provided evidence of increased brain chaoticity following propofol or xenon anesthesia, but not following ketamine anesthesia.

### 3.3 Changes in brain complexity, entropy, fractality and steepening of the spectral slope during unconsciousness are related to measures of criticality

Although the measures of criticality introduced in the previous sections are relatively new to the field of cortical electrophysiology, a wide variety of ‘classical’ EEG measures are in fact strongly related to criticality. As an example, loss of signal complexity is a widely known marker of loss of consciousness (45) but it is also characteristic of a system moving away from criticality. Thus, we sought to replicate our results on a range of criticality-related measures, which are commonly used in the field of neuroscience in a model-free manner, but whose effect may in fact be rooted in critical brain dynamics. Specifically, we applied: 1) Lempel-Ziv complexity (LZC); 2) fractal dimension; 3) multiscale entropy; 4) the Hurst exponent and; 5) the spectral slope. Analysis of the spectral slope has previously been reported for these data (32), but we included it here again to demonstrate its link to measures of criticality. All measures were estimated on 10-s windows and the full frequency range (1 – 40 Hz) and were calculated for each channel individually (see Methods). In addition, we calculated the pair correlation function (PCF) in the alpha (8-13 Hz) frequency range, which has been previously associated with ‘edge of synchrony’ criticality (6, 46) (see Discussion).

In line with previously reported results on the spectral slope (32), LZC and fractal dimension significantly decreased during propofol (LZC: *t*(4) = 6.75, *p*<0.01; fractal dimension: *t*(4) = 10.88, *p*<0.01) and xenon (LZC: *t*(4) = 7.08, *p*<0.01; fractal dimension: *t*(4) = 9.70, *p*<0.01) anesthesia, but not during ketamine anesthesia (see Fig 5A). Conversely, multiscale entropy significantly increased during propofol (*t*(4) = -9.46, *p*<0.01) and xenon (*t*(4) = -5.19, *p*<0.05) anesthesia, but not during ketamine anesthesia (see Fig 5A). The Hurst exponent captures the long-range temporal correlation of the signal and is strongly linked to both the fractal dimension and the slope of the power spectral density. Only exposure to propofol (*t(4) = -4.76, p<0.05*) yielded a significantly increased low-frequency (delta bandwidth 1-4Hz) Hurst exponent, indicating increased long-range temporal correlation during unconsciousness in the delta frequency bandwidth. In higher frequency bands, drug exposure yielded an overall decrease in Hurst exponent with alpha Hurst exponent significantly decreasing in response to xenon (t(4) = 5.55, p<0.05) (see Supplementary Material 3 for all bandwidths). In contrast to previous studies (6) the PCF did not show significant changes during general anesthesia with propofol, xenon or ketamine (see Discussion).

**Figure 5.**
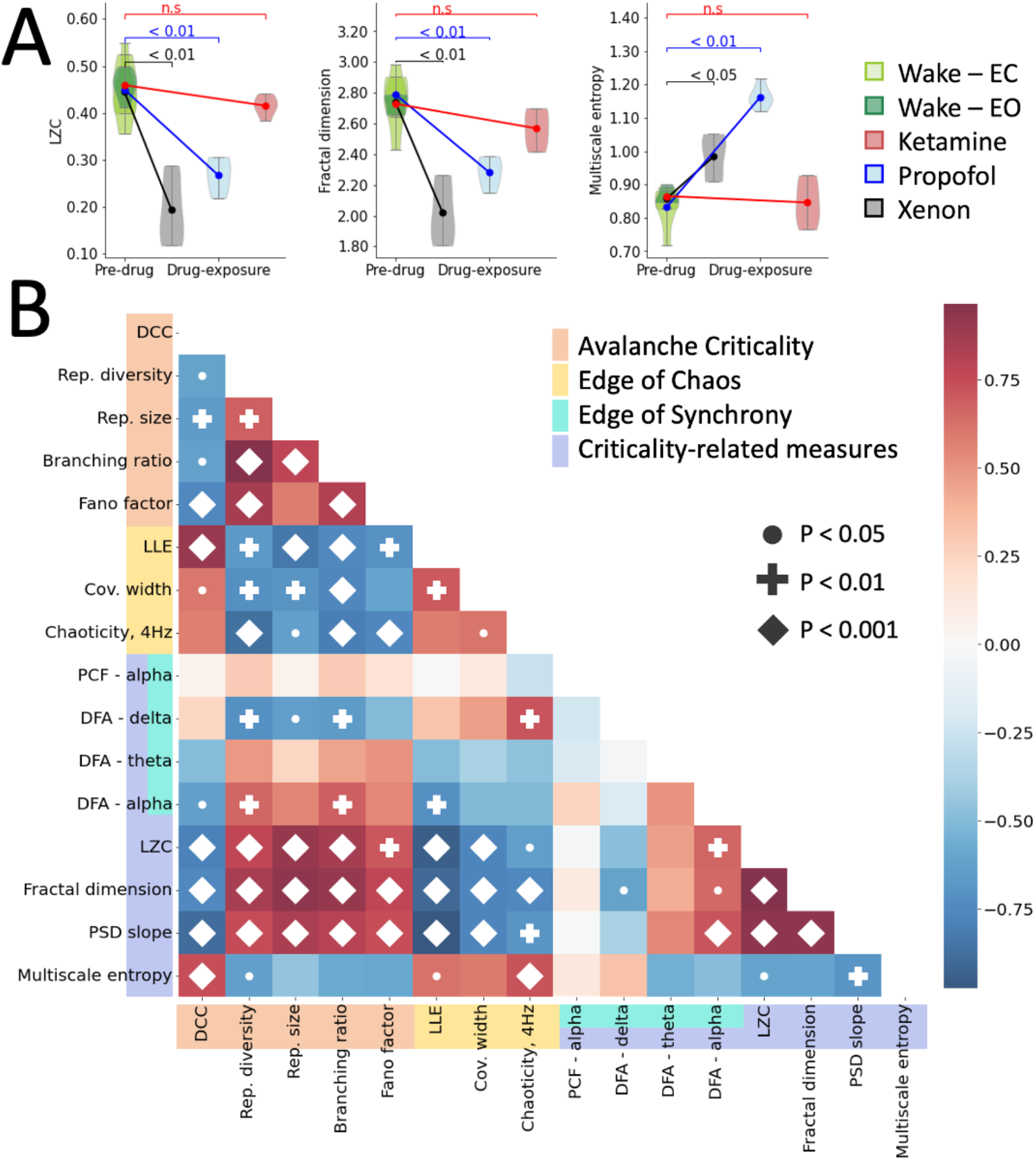
Effect of propofol, xenon and ketamine on criticality-related measures, and the mutual relation between all the applied measures. **A)** Complexity, fractal dimension and the Hurst exponent decrease during exposure to propofol (blue) and xenon (black), but not ketamine (red). Signal entropy increases during exposure to propofol and xenon, but not ketamine. Light grey lines represent individual subjects, bold lines are the mean over subjects. **B)** Correlation matrix between all criticality-related measures and measures of avalanche criticality, edge of chaos and edge of synchrony. Criticality-related measures are highly correlated with measures of avalanche criticality and edge of chaos. P-values were corrected using Bonferroni correction. Cov. width, width of the covariance distribution; DFA, detrended fluctuation analysis; DCC, deviation from criticality coefficient; LLE, largest Lyapunov exponent; LZC, Lempel-Ziv complexity; PCF, pair correlation function; PSD, power spectral density; rep., avalanche repertoire; wake - EC, wakefulness with eyes closed; wake - EO, wakefulness with eyes open.

Nearly all of these criticality-related measures are highly correlated to the above-reported measures of avalanche criticality and edge of chaos (see Fig 5B), yet no strong correlation was found with our measure of the edge of synchrony (see Discussion). This suggests that changes in brain complexity, entropy, fractality and steepening of the spectral slope observed in previous studies during unconsciousness were indicative of the brain dynamics moving away from the edge of activity propagation or the edge of chaos.

### 3.4 Criticality reliably predicts the perturbational complexity index

We next investigated the relation between the criticality of resting-state dynamics and the response to external perturbations. More specifically, we first explored the correlation between the distance from criticality of resting-state dynamics and the PCI – a measure which combines EEG and TMS to reliably detect consciousness in unresponsive patients (18). We then tested the degree of resting-state criticality as a predictor for the PCI.

Each participant’s PCI across all states (i.e., wakefulness, anesthetized) significantly correlated with all resting-state avalanche criticality measures, edge-of-chaos measures and criticality-related measures (all P < 0.01), except for the PCF (see Table 1). A direct comparison of all features between conditions is provided in Supplementary Material (see Supplementary Material 4).

**Table 1:**
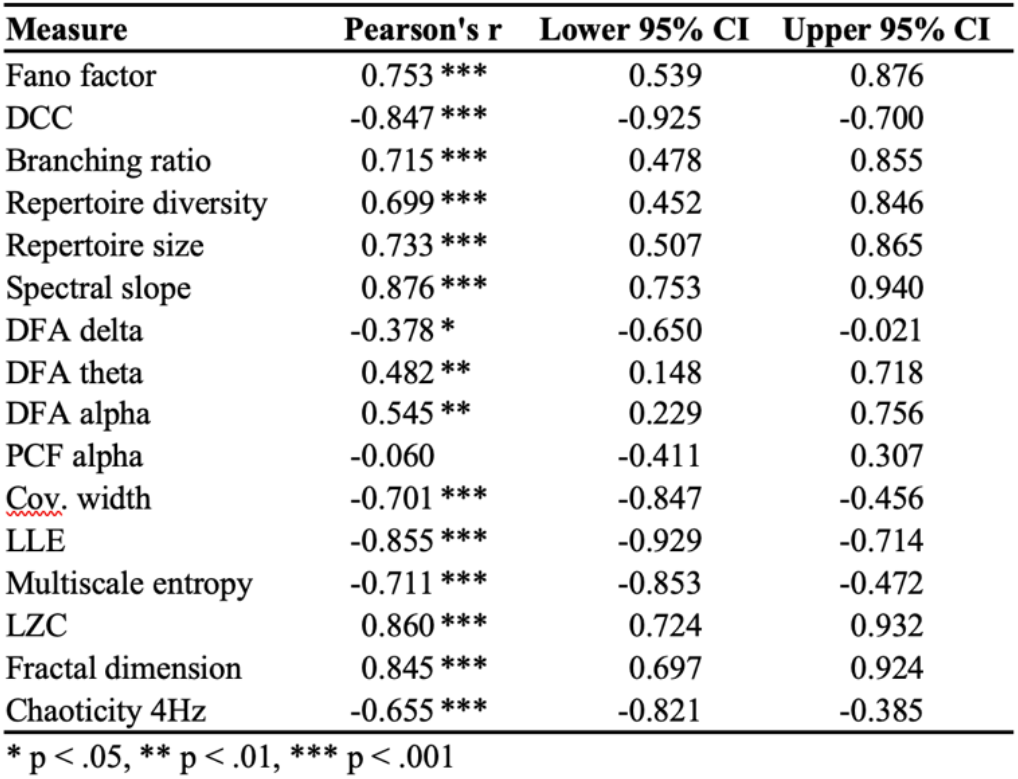
Correlation between measures of criticality and the perturbational complexity index (PCI). Cov. width, width of the covariance distribution; DFA, detrended fluctuation analysis; DCC, deviation from criticality coefficient; LLE, largest Lyapunov exponent; LZC, Lempel-Ziv complexity; PCF, pair correlation function.

| Measure | Pearson's r | Lower 95% CI | Upper 95% CI |
| --- | --- | --- | --- |
| Fano factor | 0.753 *** | 0.539 | 0.876 |
| DCC | -0.847 *** | -0.925 | -0.700 |
| Branching ratio | 0.715 *** | 0.478 | 0.855 |
| Repertoire diversity | 0.699 *** | 0.452 | 0.846 |
| Repertoire size | 0.733 *** | 0.507 | 0.865 |
| Spectral slope | 0.876 *** | 0.753 | 0.940 |
| DFA delta | -0.378 * | -0.650 | -0.021 |
| DFA theta | 0.482 ** | 0.148 | 0.718 |
| DFA alpha | 0.545 ** | 0.229 | 0.756 |
| PCF alpha | -0.060 | -0.411 | 0.307 |
| Cov. width | -0.701 *** | -0.847 | -0.456 |
| LLE | -0.855 *** | -0.929 | -0.714 |
| Multiscale entropy | -0.711 *** | -0.853 | -0.472 |
| LZC | 0.860 *** | 0.724 | 0.932 |
| Fractal dimension | 0.845 *** | 0.697 | 0.924 |
| Chaoticity 4Hz | -0.655 *** | -0.821 | -0.385 |

\* $p < .05$ , \*\* $p < .01$ , \*\*\* $p < .001$

Combining all measures in a single ridge regression model to predict PCI yielded a mean error of 0.02 (i.e., as PCI ranges from 0 to 1 this corresponds to an error of 0.2%), indicating an absolute average deviation of 0.038 ± 0.028 from the true PCI. To test the model for its predictive value on unseen data, we implemented a leave-one-subject-out (LOSO) cross-validation (n=15 splits). Model scores for each iteration were defined as the mean error of both the pre-drug and drug conditions. Using a LOSO cross-validation, the model yielded a mean error of 0.06 (i.e., 6% error), indicating an absolute average deviation of 0.067 ± 0.044 from the true PCI (see Fig. 6). Using a threshold of 0.35 yielded a perfect separation of conscious and unconscious states, for the true as well as the predicted PCI (see Fig. 6).

**Figure 6:**
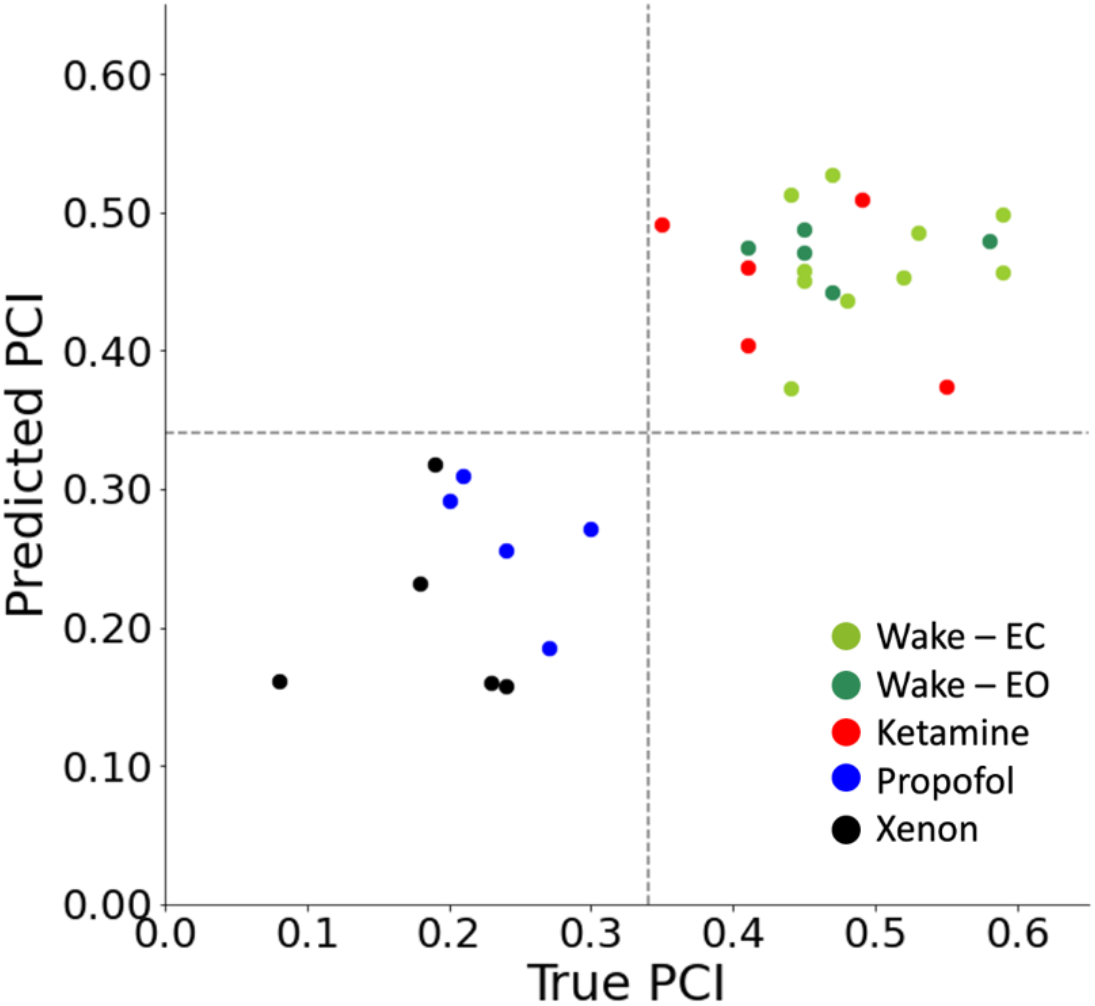
Prediction of the PCI, based on resting-state EEG dynamics. Individual points represent individual subjects during pre-drug wakefulness (light green, eyes closed; dark green, eyes open), propofol (blue), xenon (black) and ketamine (red) condition. Grey dashed lines represent a possible threshold of 0.35 to separate conscious from unconscious states.

**Figure 7:**
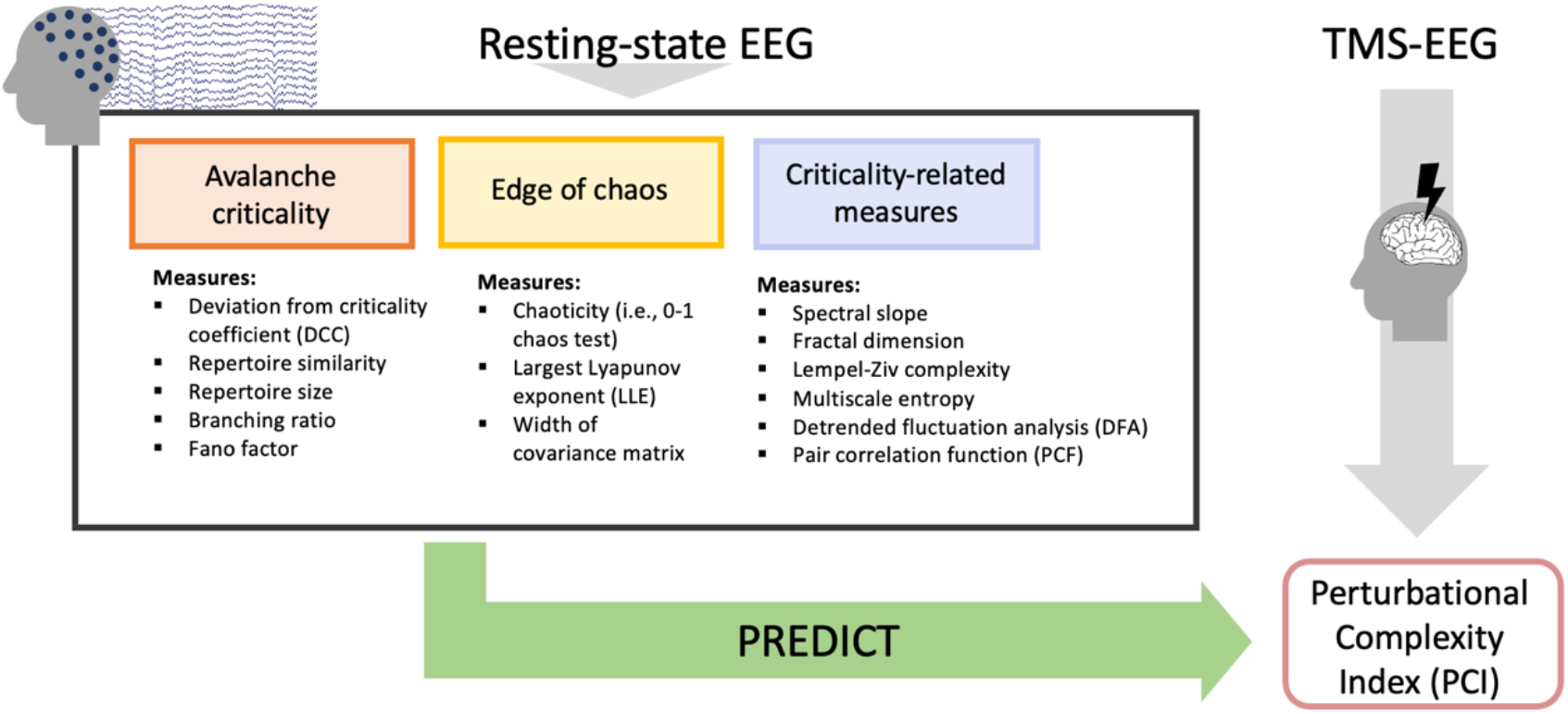
Summary of methods. We analyzed measures from two different types of criticality, avalanche criticality, edge of chaos, and a group of criticality-related measures. All measures were calculated on resting state EEG and combined to predict participants’ perturbational complexity index (PCI), as measured by EEG with transcranial magnetic stimulation (TMS).

## 4 Discussion

We investigated the effects of three anesthetics -- propofol, xenon and ketamine -- to study how different measures of brain criticality relate to the presence of consciousness beyond sheer unresponsiveness, in healthy humans. Using a variety of metrics from the classes of avalanche criticality and the edge of chaos, we observed that propofol and xenon anesthesia, which induced unconsciousness, caused brain dynamics to deviate from criticality — while ketamine anesthesia, which did not abolish consciousness, kept brain dynamics in proximity to criticality, similarly to wakefulness. We further showed that these same criticality metrics of (pre-TMS) resting-state EEG accurately predicted the brain’s PCI, supporting the notion that brain criticality may provide an explanatory framework for this reliable measure of the presence of consciousness. Together, our results provide further evidence supporting the hypothesis that critical neuronal dynamics are a necessary condition for the emergence of consciousness in the brain (3, 5, 7, 32).

In contrast to propofol and xenon, and like wakefulness, ketamine anesthesia did not abolish consciousness, nor did it result in a substantial movement away from criticality, in agreement with our hypothesis. Similar findings have been reported by Varley et al. (34) who investigated loss of complexity and critical dynamics after exposure to propofol and ketamine using invasive electrocorticography in nonhuman primates. In line with our results, propofol showed stronger loss of criticality and complexity compared to ketamine, which maintained a strong resemblance to the state of wakefulness (34). In contrast, in a human study, a sub-anesthetic infusion of ketamine (about 10% of the infusion rate used in our study) resulted in increased signal complexity (52). Similar effects have been observed using the psychedelic compound lysergic acid diethylamide (LSD), which increased the complexity (7, 52), and reduced the chaoticity of the EEG signal, thus narrowing its distance from the edge of chaos (7). This highlights the importance of differentiating between sub-anesthetic and anesthetic doses of ketamine, which have distinct effects on brain complexity and criticality.

Beyond providing an explanatory mechanism for the PCI, our results may also have major practical implications for the clinical assessment of human consciousness. PCI is a well-characterized metric that reliably distinguishes conscious from unconscious states (15, 18), yet the requirement of a TMS system and a long testing procedure has limited its wider application in clinical practice. In this study, we demonstrated the relationship between EEG criticality and the PCI by predicting individual subjects’ PCI using only short resting-state 60-channel EEG recordings obtained just prior to TMS intervention, with a mean average error below 7%. Although our results suggest that the assessment of consciousness could be accurately carried out without requiring TMS, we highlight the complementarity of these approaches. While the complexity of the brain’s response to TMS can be explained through network criticality, measuring the PCI yields more information than its single summary complexity value PCI_max_. Specifically, each TMS stimulation in the PCI procedure produces a detailed cortical map with spatial information about which regions were or were not affected by the perturbation (18, 53). Especially for brain-injured patients (53), this information contains meaningful clinical insights above and beyond the final index that cannot be predicted based on resting-state criticality alone. Whether or not one could in a clinical context rely on criticality measures to derive an estimate of PCI without the need to actually stimulate the brain, as was found here in healthy controls undergoing anesthesia, still remains to be seen.

The association between criticality and the complexity of evoked responses has previously been demonstrated in cortical cultures (23, 24) and in silico modeling (50). Shew et al. (23, 24) measured stimulus-evoked pattern entropy during drug-induced over-excitation and -inhibition and demonstrated that the diversity of patterns was maximized at the critical balance between excitation and inhibition, where neuronal avalanches obeyed scale-free distributions. Aligning with the results of the present study, Shew et al. (23) concluded that “spontaneous activity and input processing are unified in the context of critical phenomena”. In addition, Momi, Wang and Griffiths (51) used whole-brain connectome-based computational modelling to reconstruct individual responses to TMS in vitro and highlighted the role of GABAergic neural populations and cortical excitability. Our results align with this previous research (23, 24, 49–51) and support the hypothesis that the complexity of the response following perturbation can be inferred solely based on resting-state activity.

In this study, we focused our analysis on measures of two types of criticality, namely, avalanche criticality and the edge of chaos (for detailed discussion on the types of criticality and their relation to consciousness see Supplementary Methods, Note1). While both types of critical phase transitions are theoretically distinct and dissociable (41, 42, 54), deviations from these respective critical points may nonetheless be correlated in specific classes of systems (Haldeman & Beggs, 2005). More work will be needed to understand their interrelation in the brain. Other types of criticality have also been studied in brain networks, including the “edge of synchrony” (see 2 as a review).

For avalanche criticality, propofol- and xenon-induced unconsciousness shifted the brain network away from criticality and yielded subcritical dynamics. While the shift away from criticality aligns with our hypothesis, we tentatively expected subcritical dynamics only during propofol anesthesia (over-inhibition yielding a reduced spread of perturbations), but supercritical dynamics during xenon anesthesia (over-excitation yielding strong but uniform reactions to perturbation). Instead, we observed a shift towards subcritical dynamics for both anesthetic conditions. Similar results were observed by Colombo et al. (32), where exposure to both propofol and xenon anesthesia yielded the same electrophysiological effect, namely an overall slowing of the EEG and steepening of the spectral slope. Indeed, xenon functions as a N-methyl-D-aspartate (NMDA) antagonist, reducing NMDA-activated currents by about 60% (55, 56). However, xenon has also been proposed to yield unconsciousness due to over-excitation (57). The strong but stereotypical response to perturbation using TMS could result from a possible state of neural bistability induced by xenon (15, 32), in which the oscillation between strong depolarization and hyperpolarization could provoke high-amplitude EEG, despite overall subcritical dynamics. It is interesting to note here that despite larger-amplitude signal fluctuations under propofol and xenon, there were nevertheless fewer and smaller avalanches observed in these conditions. This highlights the dissociation between signal power and avalanche dynamics.

In terms of chaos, exposure to propofol or xenon, but not ketamine, yielded an increase in chaoticity with respect to normal wakefulness. What does this say about the relationship between consciousness and the edge of chaos? Canonically, a positive largest Lyapunov exponent (LLE) indicates the presence of chaos, with the edge of chaos situated at LLE = 0. For the 0-1 chaos test, the K-median value corresponding to the edge of chaos is less clear, but previous work using similar methods and a ~4 Hz low-pass situated this value around K-median = 0.85 (7) The present results, with positive LLE and K-median ≅ 0.85 would then indicate that the neural dynamics of waking consciousness operate near the edge of chaos, slightly in the chaotic phase, and that the unconsciousness induced by xenon and propofol exposure is accompanied by a shift of the dynamical operating point *away* from edge-of-chaos criticality and further into the chaotic phase. Meanwhile, neural dynamics under ketamine exposure remained indistinguishable from waking dynamics, remaining close to the edge of chaos. For a discussion of the significance of finding waking neurotypical dynamics slightly away from the critical point, see O’Byrne & Jerbi (2).

In addition to metrics that were derived from criticality theory, our study also examined ‘criticality-related’ metrics; in other words, metrics that are: (i) commonly used in electrophysiology and; (ii) expected to bear a strong relation with the distance from criticality. These predominantly showed strong correlations with the theoretically derived metrics and demonstrated the expected relationships with consciousness: LZC and fractal dimension each decreased with loss of consciousness. However, MSE unexpectedly increased with unconsciousness; furthermore, it was less strongly correlated with the theoretically derived metrics. The relationship between multiscale entropy and criticality is still unclear, with some recent work indicating that such measures of randomness continue to increase past the critical point and into the supercritical or chaotic phase (58). Indeed, MSE in our data was strongly correlated with chaoticity.

We were not able to replicate previous findings of reduced alpha-band PCF during anesthetic-induced unconsciousness (6, 59). In addition, the PCF did not correlate with participants’ level of consciousness, as measured by the PCI. However, since we were particularly interested in computationally simple measures for clinical applicability, the PCF in this study was estimated on sensor-level and not source-localized EEG, in contrast to previous studies (6, 59). In addition, synchrony-based measures of criticality rely on the presence of narrowband oscillations, which can pose methodological or conceptual challenges given the strong spectral changes usually observed during pharmacologically-induced unconsciousness (32) or the total absence of oscillatory peaks in some forms of pathological unconsciousness (60, 61).

The results of the present study need to be considered in the light of some limitations. First, this study was conducted on a dataset of only 15 healthy adults. It should be the subject of future research to replicate these results on a larger cohort and across a wider variety of pharmacological, pathological and physiological states of unconsciousness.

Second, this study explored a range of measures from the categories of avalanche criticality, edge of chaos and criticality-related measures; however, this battery of measures is by no means exhaustive and was selected based on translatability to scalp-level human EEG. This study does not aim to promote a specific set of measures, but rather to motivate further exploration of the framework of criticality as a requirement for human consciousness.

Third, in our data, the distribution of avalanche sizes followed a truncated power law (i.e., a power law with an exponential tail at large scales) in 25 out of 30 recordings. The fact that most data exhibited truncated power laws - instead of fully scale-free power law behavior - can be attributed to the finite size of the system (so-called finite-size effects) and to the relatively small amount of data, which limits the possibility of observing large avalanches, yielding an exponential drop-off at larger scales. The result that lognormal distributions yielded better fits in 5 subjects can be attributed to more extreme cases of the same causes (37).

Fourth, the distance to edge of chaos is difficult to quantify with certainty in our data. As noted above, the 0-1 chaos test used in this study does not directly provide an estimate of this distance. Likewise, while the width of the covariance matrix can theoretically be used to precisely measure the distance to criticality in multi-unit recordings (41), it is less clear how to do so in coarser-grained recordings like EEG. The LLE as measured here using Rosenstein’s method (44) is our most straightforward indicator of the distance to criticality, with LLE = 0 indexing the edge of chaos, but further validation of this measure in brain recordings will be needed.

Fifth, previous studies have shown a direct link between the distance to criticality of a network’s spontaneous activity and the complexity of the network’s reaction to perturbations (23, 24). However, Shew et al. (23, 24) measured the response to perturbations immediately following the measurement of resting-state dynamics. In contrast, the PCI requires repeated stimulation over a period of several minutes and subsequent averaging of recorded effects. The time delay between the recorded resting-state EEG and the end of the TMS protocol required to obtain the PCI might be a source of variability, especially for patients with disorders of consciousness, where levels of consciousness and wakefulness quickly fluctuate over time.

Lastly, the present study only draws a relation between resting-state network criticality and PCI for the assessment of *pharmacologically induced* unconsciousness, which does not allow generalization to *pathological* loss of consciousness. Patients with disorders of consciousness were previously found to exhibit cortical dynamics far from the edge of chaos, with dynamics approaching the edge of chaos upon recovery (7). However, Liu et al. (62) highlighted a stark difference in scale-free properties of functional network interactions between patients in a minimally conscious state and anesthetized healthy adults. It is therefore too early to conclude whether criticality can be used to reliably assess consciousness in clinical populations with damaged brain network integrity.

In summary, this study demonstrates that propofol- and xenon-induced unconsciousness is accompanied by a distancing from criticality, as measured by avalanche criticality, chaoticity and criticality-related measures. In contrast, ketamine anesthesia did not significantly alter the distance from criticality, remaining indistinguishable from wakefulness in dynamical space. Furthermore, using the dynamical properties of resting-state EEG only, we were able to predict the PCI with a mean error below 7%, without the use of a TMS machine. Criticality can be seen as a unifying framework which binds concepts of complexity, integrated information and sensitivity to perturbation into a coherent narrative. This study supports the hypothesis that critical brain dynamics are implicated in the emergence of consciousness and may provide new directions for the clinical assessment of consciousness.

## 5 Materials and methods

### 5.1 Participants and anesthetic protocol

We analyzed 15 healthy subjects (5 males, 18–28 years old) from an existing dataset, previously published by Sarasso et al. (15). Each of the 15 subjects provided written informed consent and was randomly assigned to a group whereupon they were exposed to general anesthesia either with propofol (n=5), xenon (n=5) or ketamine (n=5), in absence of any surgical procedure. The study was approved by the ethical committee of the University of Liège (Liège, Belgium). EEG data were recorded using a TMS-compatible 60-channel EEG amplifier (Nexstim Plc., Finland). Before the start of the anesthetic protocol, 10 min of resting-state EEG were recorded, followed by a 6 to 8 minute-long protocol of TMS-EEG (15). Whereas the stimulation of different cortical targets can yield different values of PCI (18), the present study only considers the maximum among these values (PCI_max_), which is the value typically used to evaluate the presence of consciousness. During the TMS-EEG protocol, up to 250 stimuli were conducted over a single stimulation site (Brodmann Area 6 or 7) (15). Each of the three anesthetic procedures (propofol, xenon, ketamine) aimed at reaching a common behavioral state of unresponsiveness, i.e., a Ramsey Scale score of 6, following systematic repeated assessments, corresponding to a ‘surgical level’ of anesthesia. Propofol was administered through a target-controlled infusion pump (Alaris TIVA; CareFusion), using a target-effect concentration of 3 μg/ml. Xenon was administered by inhalation (62.5 ± 2.5 % in oxygen). Ketamine was administered through a 2 mg/kg intravenous infusion (see 15 for full details).

After the target concentration was reached, continuous EEG was acquired for a period of 3 to 5 minutes before the TMS-EEG protocol. Upon awakening from behavioral unresponsiveness, retrospective reports were collected from each participant as a proxy for presence or absence of consciousness (see 15 for full details).

### 5.2 Electroencephalography data

The 60-channel resting-state EEG data were preprocessed for a previously published study (32). In brief, the signal was filtered between 0.5 and 50 Hz, bad channels were rejected by a trained experimenter, and rejected channels were interpolated by spherical splines. Data segments with excessive levels of noise were manually rejected. Independent component analysis was performed to reduce muscle and eye movement artifacts. A minimum of 1.5 minutes and a maximum of 5 minutes of clean resting-state EEG data were selected for analysis (265 s ± 64s).

### 5.3 Avalanche criticality analysis

Many dynamical systems away from equilibrium exhibit a typical behavior of so-called “avalanches” – chains or cascades of activity that propagate across the network (space) and across time. At criticality, these avalanches become generically scale-invariant; that is, the probability distributions of various avalanche properties follow a power law. In the brain, ‘neuronal avalanches’ are measured by a thresholding and binning of the electrophysiological time series, a method first developed by Beggs and Plenz (28). This method depends on two hyperparameters: the signal binarization threshold (or event detection threshold) and the time bin.

The binarization threshold was determined using a data-driven method (33, 34), whereby the EEG signal was first z-transformed channelwise (by subtracting each value by the signal mean and dividing by the SD) and plotted in a probability distribution ranging from –10 to 10 SD. These probability distributions were then averaged across all participants within a same condition (Fig. 2B). A Gaussian was fit to each of these distributions, and the binarization threshold was taken as the point of divergence of the data distribution from the Gaussian, with the rationale that a divergence from the Gaussian reveals signal that is unlikely to arise from mere stochastic fluctuation.

The signal was binarized by identifying signal excursions above (below) the positive (negative) bthreshold, and for each excursion, the maximum (minimum) value of the excursion was set to one, and all other values set to zero. Avalanches were then identified by scanning forward in time through the multichannel binarized data and finding a first neural event (a one among the zeros), then looking for additional events (on any channel) occurring within a delay less than or equal to the time bin. If an event is found, it is added to the avalanche and the process is iterated again. If no events are found within the time bin, the avalanche ends, and the process is begun again at the next occurrence of an event to find the next avalanche. These methods are standard practice for the detection of neuronal avalanches (28, 33). Fitting and maximum likelihood estimation for power laws and other functional forms was carried out using the *powerlaw* Python package (37). Other avalanche criticality analyses were carried out using the *edgeofpy* Python package, available at https://github.com/jnobyrne/edgeofpy.

#### 5.3.1 Deviation from criticality coefficient

Neuronal avalanches are described mainly by their size S (in EEG, the number of contributing electrodes), the avalanche duration T, and the average S for every T. The exponents of the power laws of these distributions are known as critical exponents and are referred to by τ, α and 1/σ*vz*, respectively (28, 29)

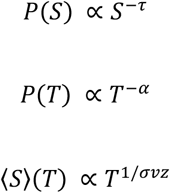

Certain predictions exist for the values of these exponents in certain classes of systems (29, 35); however, it is still unclear what these values should be in the brain (63). Still, whatever the values, it is expected that the exponents of systems at criticality (for a broad range of universality classes) should obey the following scaling relation (29, 35):

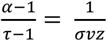

The degree to which the above relation is followed by the neuronal avalanche data is a good indicator of the brain’s proximity to avalanche criticality. We therefore define the deviation from criticality coefficient (DCC) as (38):

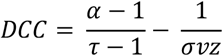

#### 5.3.2 Branching parameter

The branching parameter *m* (also called branching ratio) is a measure of activity propagation, describing the average number of events resulting as descendants from one single event. Critical systems are characterized by a branching parameter of *m* = 1 (i.e., one event is followed on average by exactly one event), enabling activity to be stably propagated through the system. Subcritical systems exhibit values *m* < 1 (i.e., one event is on average followed by less than one event), resulting in a typically fast vanishing of activity. In contrast, supercritical systems are characterized by *m* > 1 (i.e., one event is on average followed by more than one event), resulting in a fast amplification of activity. The branching parameter was defined as the number of events in time bin t divided by the number of events in the preceding time bin t-1, averaged over all time bins t (28). As the branching ratio is highly sensitive to the chosen length of the time bin, all results were replicated using a range of time bins from 1 ms to 12 ms.

#### 5.3.3 Avalanche repertoire size and similarity

As neuronal avalanches spread throughout the cortex, they exhibit a variety of spatial patterns (i.e. the combination of electrodes activated during a given avalanche). The size of the avalanche repertoire was defined as the number of unique avalanche patterns (40) and was normalized by the length of the signal. Whereas large values indicate a wider range of different activation patterns, small values indicate that activity is driven by a smaller number of repeating patterns. Avalanche repertoire diversity was estimated using the median normalized Hamming distance between all identified unique patterns (40). Low repertoire diversity values indicate high similarity between existing patterns, and large values indicate highly dissimilar patterns.

#### 5.3.4 Fano factor

The Fano factor (FF) is a measure of the variability of a signal, and is expected to peak for critical processes (65, 66) It is defined as:

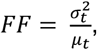

Where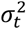 and µ_*t*_ are the variance and mean of the signal over time t, respectively.

### 5.4 Edge of chaos analysis

#### 5.4.1 Modified 0-1 chaos test

Signal chaoticity (K) was estimated using the modified 0-1 chaos test (43, 67). The signal was epoched on non-overlapping 10-s windows. K was estimated using a Python translation of the code provided by Toker et al. (7) (available in *edgeofpy*). It was calculated on every channel and non-overlapping 10 s epoch individually and averaged over time. Chaoticity of the whole brain network was defined as the median K over all electrodes. The use of K-median for cortical dynamics has only been validated on slow cortical dynamics (7). Therefore, the signal was low-pass filtered at a range from 1 to 12 Hz prior to the estimation of chaoticity. In a second approach, we used the FOOOF algorithm (68) to identify oscillatory peaks between 1 and 6 Hz for every channel and epoch individually. A low-pass filter set to the maximum frequency of the oscillatory peak was applied to the corresponding channel segment. Channels without an oscillatory peak were excluded from the FOOOF-based chaoticity analysis. These results are reported in Supplementary Material 2.

#### 5.4.2 Largest Lyapunov exponent

The Lyapunov exponent (λ) is a measure of sensitivity to initial conditions and estimates how much the trajectories of two initially neighbouring points converge or diverge over time. λ was calculated using the Neurokit2 implementation (69) of the Rosenstein method (44). Whereas values of λ < 0 indicate stable dynamics (i.e. trajectories converge over time), values of λ > 0 indicate chaotic dynamics (i.e. initially close trajectories diverge over time). The estimation of λ requires reconstruction of the signal state space, which was created using delay-embedding (delay=1, dimension=2). Closest neighbours were detected based on Euclidean distance. A least-square fit was then used to estimate the slope (i.e. λ) of the distance line. λ was calculated on every channel and non-overlapping 10-s epoch individually and averaged subsequently over time.

#### 5.4.3 Width of covariance matrix

In neural networks, the onset of chaos occurs when the spectral radius, i.e. the largest eigenvalue of the effective connectivity matrix, λ_ECM_, is larger than one, indicating the presence of an unstable (chaotic) eigenmode. In general, the brain’s effective connectivity matrix is difficult to estimate from brain recordings, especially under subsampling. However, according to analytical work by Dahmen et al. (41), the integrated time-lagged covariance matrix estimated from subsampled recordings can provide unbiased information about the largest eigenvalue of the underlying effective connectivity matrix λ_ECM_. Specifically, the normalized width of the distribution of covariances Δ is positively and monotonically related to λ_ECM_, and thus to the degree of instability or chaos in the network dynamics. Here, we first calculated the integrated time-lagged covariance matrix (also known as noise covariance), then estimated Δ as the standard deviation of the off-diagonal elements of the covariance matrix divided by the mean of the diagonal elements. For details on these analyses, see Dahmen et al. (41) and Morales et al. (70). This analysis was carried out using the *edgeofpy* Python package.

### 5.5 Criticality-related measures

#### 5.5.1 Detrended fluctuation analysis

The Hurst exponent was calculated using the Neurokit2 implementation (69) of DFA. Due to variable available signal lengths, we used a maximum of 200s of data for the DFA analysis. DFA was calculated on each channel individually (using all available data per channel) and on a range of scales from 1 to 20 s (as recommended for this method, the upper limit of the range being one tenth of the signal length). DFA was performed on the amplitude envelope of the delta, theta, alpha, beta and gamma bands individually, as described by (71). The amplitude envelope was extracted using the absolute value of the signal’s Hilbert transform. Hurst exponents were calculated on every epoch and channel individually and averaged over time.

#### 5.5.2 Spectral slope

The spectral slope, or aperiodic slope, describes the decay of the power spectral density (PSD) (i.e., the exponential decay of power over frequency) (72). The PSD was estimated using the Welch method for every 10-s epoch and channel individually. The spectral slope was estimated epoch-wise using the FOOOF package over a frequency range of 1 to 40 Hz (68) and averaged subsequently.

#### 5.5.3 Complexity

Univariate Lempel-Ziv complexity (LZC) was estimated using Neurokit2 (69). LZC was calculated on every channel and non-overlapping epoch of 10s independently and averaged subsequently. The signal was binarized using the mean of each channel and epoch individually. LZC was normalized using the length of the sequence (73).

#### 5.5.4 Multiscale entropy

Multiscale entropy was estimated using Neurokit2 (69). Multiscale entropy calculates sample entropy on several timescales using a coarse-graining approach (74, 75). The optimal embedding dimension for the entropy estimation was calculated using average false nearest-neighbors method implemented in Neurokit2 (69). Multiscale entropy was defined as the sum of sample entropy values over all scales (69).

#### 5.5.5 Fractal dimension

Fractal dimension was estimated using the Neurokit2 implementation (69) of Katz’s fractal dimension. While other methods are available for the estimation of fractal dimension, this algorithm has been shown to be more robust against noise (75). The Katz algorithm for fractal dimension estimates the sum of Euclidean distances between all successive signal points and identifies the maximum distance between any starting point and any other point in the signal (69).

#### 5.5.6 Pair correlation function

The pair correlation function (PCF) is a measure of the variance of phase-coupling in a system of oscillators, with higher values indicating a higher susceptibility and closeness to critical dynamics (46). PCF was estimated using a custom Python function (available in *edgeofpy*). Prior to calculation, the signal was downsampled to 250 Hz and bandpass-filtered in the alpha frequency range (8-13 Hz). The PCF was estimated on every non-overlapping 10-s epoch individually and subsequently averaged over time.

### 5.6 Statistical analysis

The difference between metrics derived during wakefulness and metrics derived during xenon, propofol or ketamine anesthesia was assessed using a repeated-measures t-test for each group individually. P-values were corrected for multiple comparisons using the Holm correction. For statistical tests on topographically distributed channels and the visualization of significantly changing brain regions, p-values were corrected using permutation cluster tests. Correlation to the PCI was assessed using Pearson correlation. For the prediction of the PCI, a multivariate ridge regression (*alpha* = 1) was trained on 14 features (i.e., DCC, repertoire diversity, repertoire size, branching ratio, Fano factor, LLE, width of the covariance matrix, chaoticity estimate at 4 Hz low-pass filter, alpha-band PCF, DFA, LZC, fractal dimension, spectral slope and multiscale entropy) to predict the PCI_max_ value for each patient and condition. To test the model’s predictability, we implemented a leave-one-subject-out cross validation (i.e. 15 folds, one for each subject). The model was trained 15 times on 14 subjects and tested on both conditions of the corresponding hold-out subject. The mean error was defined as the average absolute difference between the predicted and the real PCI_max_ values over all conditions and subjects.

## 6 Acknowledgments

CM is funded through Fonds de recherche du Québec – Santé (FRQS). JOB is supported by the Canadian Institutes of Health Research (CIHR) and FRQS. OG is research associate and SL is research director at F.R.S-FNRS. SL is Research Director at the Belgian National Fund for Scientific Research, Chairholder of the Canada Excellence Research Chair in Integrative Neuroscience for Sustainable Mental Health and funded by European Foundation of Biomedical Research FERB Onlus. K.J. was supported by funding from the Canada Research Chairs Program and a Discovery Grant (grant no. RGPIN-2015-04854) from the Natural Sciences and Engineering Research Council of Canada (NSERC), a New Investigators Award from the Fonds de recherche du Québec en nature et technologies (FRQNT) (grant no. 2018-NC-206005), and an Institut de valorisation des données (IVADO) fundamental research project grant funded through the Canada First Research Excellence Fund (CFREF) program. SBM is supported by the Canada Research Chairs Program (Tier II). This research was funded through the FRQNT Strategic Clusters Program (2020-RS4-265502 - Centre UNIQUE - Union Neurosciences & Artificial Intelligence – Quebec, an NSERC Discovery Grant (RGPIN-201603817), the Canada Excellence Research Chairs Program (#215063), the Canadian Institutes of Health Research (#408004), the ERA-Net FLAG-ERA JTC2021 project ModelDXConsciousness (Human Brain Project Partnering Project) and the NINDS grant 1K23NS112473. This research was undertaken thanks in part to funding from the Canada First Research Excellence Fund and Fonds de recherche du Québec, awarded to the Healthy Brains, Healthy Lives initiative at McGill University, and the International Anesthesia Research Society (IARS)

