## Supplementary Material for "Criticality of resting-state EEG predicts perturbational complexity and level of consciousness during anesthesia"

**This document contains:**


Supplementary Material, Figure 1

Supplementary Material, Figure 2

Supplementary Material, Figure 3

Supplementary Material, Figure 4

Supplementary Methods

- Note 1
- Note 2

### Supplementary Material 1

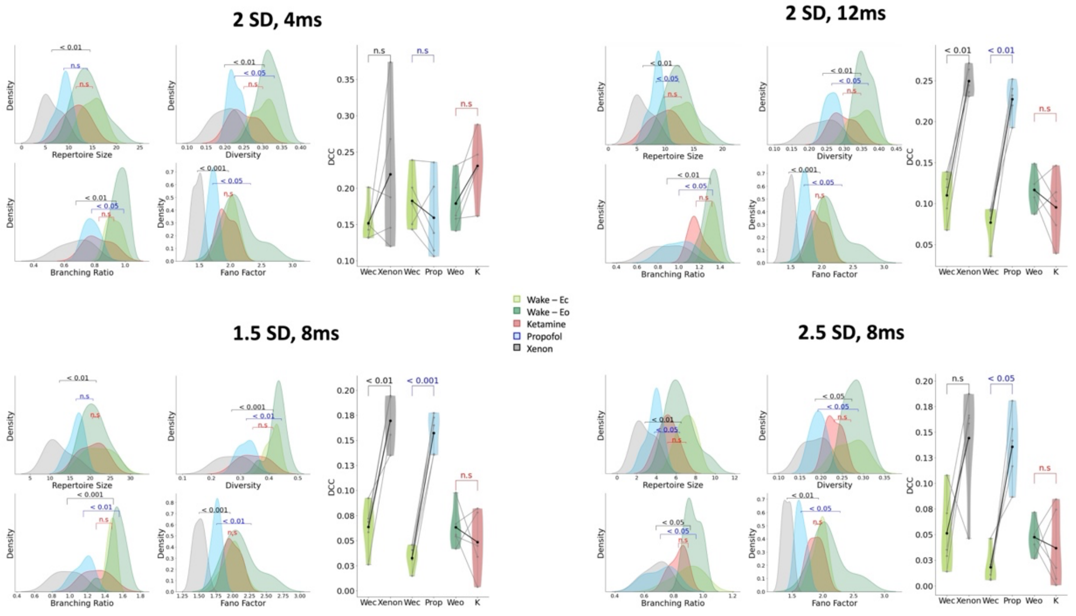


**Supplementary Figure 1:** From the binarized signal, avalanches were detected using an inter-event interval of 4-12 ms and using 1.5 to 3 SD. All results reported in the main manuscript (using 8 ms and 2 SD) were replicated for a range of 1.5 to 3 SD (lower panels) and for an increased inter-event interval of 12 s (top right panel). Decreasing the inter-event interval to 4 s only allowed a partial reproduction of the original results (top left panel). As reported in the main manuscript, the deviation from criticality coefficient (DCC) increases under the effect of xenon and propofol, but not ketamine. Light grey lines represent individual subjects, bold lines are the mean over subjects. Effect of xenon, propofol and ketamine on measures of criticality: xenon and propofol but not ketamine resulted in decreases in the size and diversity of the avalanche pattern repertoire, the branching ratio and the Fano factor, each of which is expected for more subcritical dynamics.

### Supplementary Material 2

The median K-statistic was estimated using the modified 01 chaos test (1) on the low-pass filtered signal of each channel and epoch individually. Whereas the main manuscript contains values of chaoticity using a range of fixed low-pass filter frequencies, chaos analysis was repeated using a modular low-pass frequency, as proposed by Toker et al. (2). The lowest oscillatory peak on every epoch and channel was estimated using the FOOOF algorithm (3). Data was low-pass filtered using the higher edge of the identified oscillation between 1 to 6 Hz. If no oscillatory peak could be detected, the channel in the particular segment was rejected from further analysis.

The absence of oscillatory peaks and thus exclusion of the corresponding data segment concerned 17 % of the data during wakefulness and propofol anesthesia, 20 % during xenon-induced unconsciousness and 45 % after the administration of ketamine. For the remaining data, an average low-pass frequency of 3.53 ± 0.29 was selected (see Supplementary Figure 2).

Using a flexible filtering frequency, signal chaoticity significantly increased after administration of propofol anesthesia (*P* < 0.05), but not in response to xenon or ketamine. The high variability in low-pass frequencies, as well as the overall absence of low frequency oscillatory peaks during exposure to ketamine demonstrates the limitation of this method for the comparison of groups with varying spectral properties.

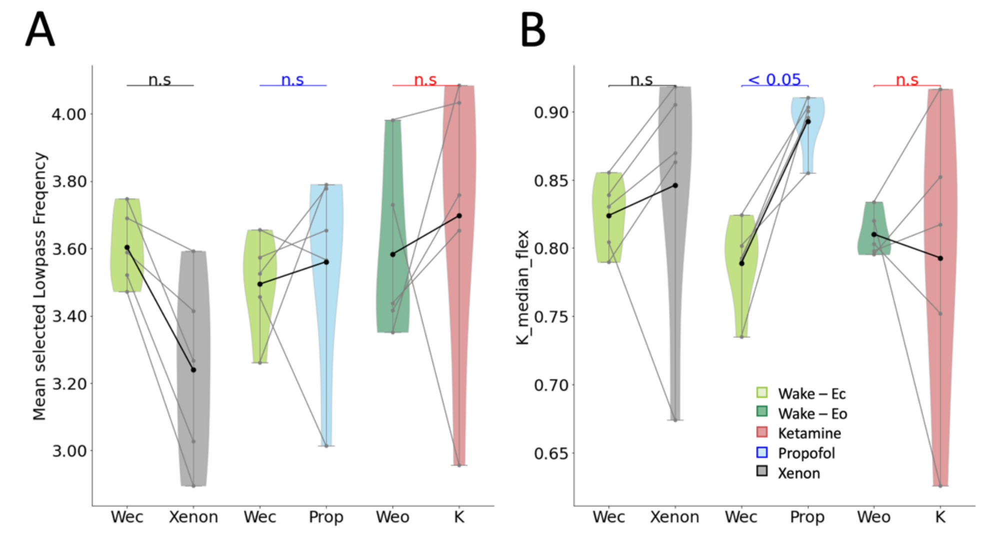


**Supplementary Figure 2:** Effect of exposure to propofol, xenon and ketamine on K-median using a variable low-pass frequency. **A)** Average selected low-pass frequency using the FOOOF algorithm, **B)** K-median using a variable filter frequency. Chaoticity significantly increases only during exposure to propofol. Light grey lines represent individual subjects, bold lines are the mean over subjects.

### Supplementary Material 3

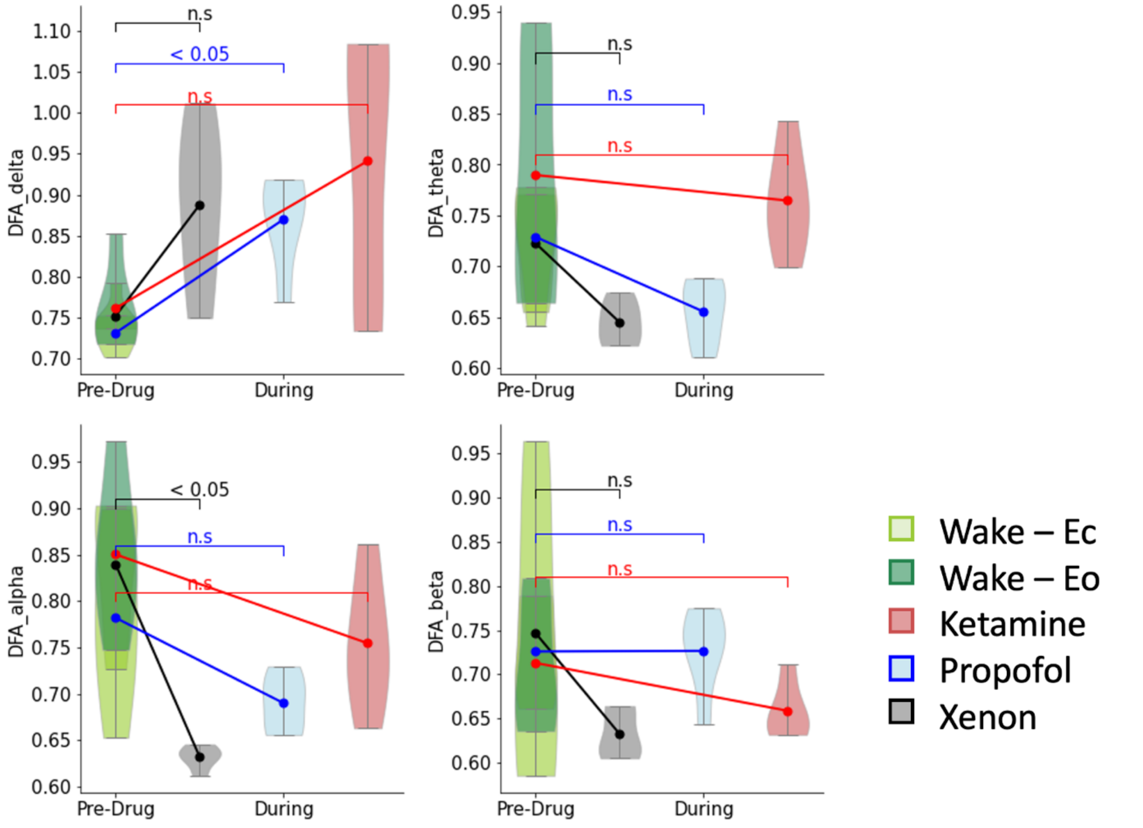


**Supplementary Figure 3:** Effect of exposure to propofol, xenon and ketamine on Detrended Fluctuation Analysis Hurst component in different frequency ranges. P-values not corrected for multiple comparison of frequency bands.

### Supplementary Material 4

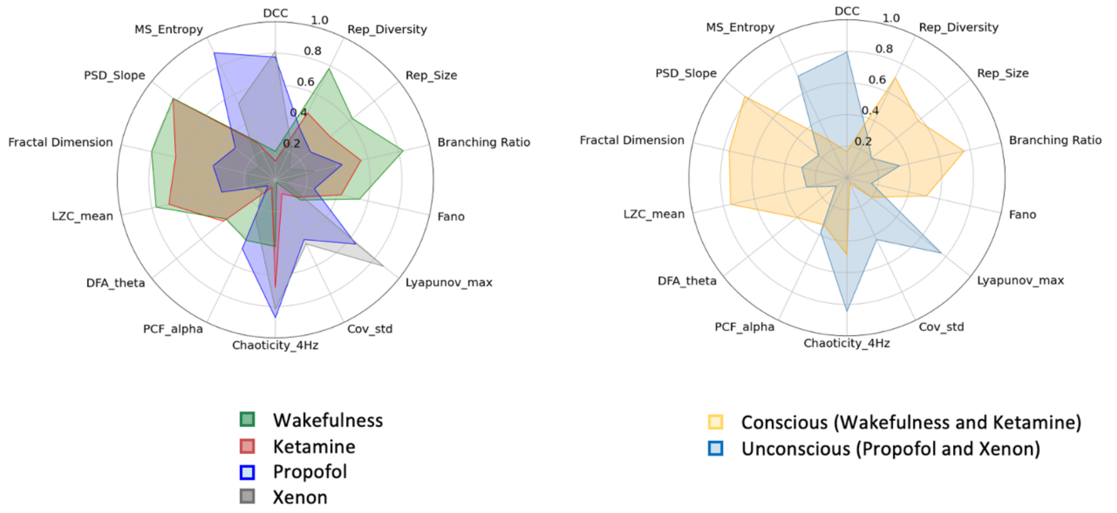


**Supplementary Figure 4:** Comparison or relative change of measures of criticality in wakefulness and drug exposure (left) and conscious and unconscious states (right). All features were normalized on a scale from 0 to 1 only for the purpose of visualization and should not be interpreted individually.

### Supplementary Methods, Note1

*Avalanche criticality and consciousness*

Avalanche criticality describes a class of critical (second-order) phase transitions where dynamics switch from amplification to dissipation of activity, i.e. where in one phase, single-node perturbations to the network tend to cascade and 'snowball' until the whole network is activated (amplification), while in the other phase, perturbations cascade very little, if at all (dissipation). At the critical point, there is no characteristic scale to these cascades, such that a single-node perturbation can activate small, intermediate or system-size portions of the network. With regard to consciousness, the amplification and dissipation phases bear a close correspondence with concepts of integration and differentiation, as follows. A system far into the amplification (supercritical) phase will be highly integrated, in the sense that activation of any single node will be propagated (in the energetic sense)—or communicated (in the informational sense)—to every other node with high likelihood; however, this also means that the dynamics are poorly differentiated, in that every possible single-node input tends to yield the same system-wide activation response. Moreover, the long lifetime of perturbations in the supercritical phase entails that inputs presented close together in time will interfere, and their information will become corrupted. Conversely, a system far in the dissipation (subcritical) phase will show highly differentiated (information-rich) input-output relations, but will do so in a disintegrated manner, whereby the activation of one node has little or no impact on the activity of most other nodes. The meeting point of these two phases at the critical point harnesses the best of both worlds, where large, integrating avalanches coexist with small differentiating avalanches. Therefore, by the intuition that a balance of integration and differentiation should be necessary for consciousness, avalanche-critical brain dynamics are an excellent candidate mechanism for this balance supporting consciousness. Indeed, some prior work supports this hypothesis (5–8). In line with this work, we here showed, through a variety of metrics, that conscious states displayed electrophysiological signatures closer to avalanche criticality than anesthetic-induced unconscious states.

*The edge of chaos and consciousness*

The edge of chaos has been less extensively studied in the brain, with the concept taking its roots in computer science and dynamical systems theory (9, 13, 14). It is the type of critical point that separates a chaotic phase from a stable phase in a system's dynamical phase diagram. In the stable phase, small differences in the system's initial conditions are 'forgotten' as the system quickly evolves towards fixed-point or periodic attractors in its state space, while in the chaotic phase, small differences are blown up exponentially with time, leading to vast differences in outcomes stemming from minute differences in inputs. In finite-dimensional systems, this equally leads to 'forgetfulness', as the effect of past inputs is rapidly corrupted by the explosive diffusion of new inputs. At the transition between stable and chaotic phases—which has been dubbed the "edge of chaos" but may just as well be called the "edge of stability" or "edge of instability"—small differences are neither overlooked nor overblown but are faithfully and persistently maintained in the system dynamics. This confers the system with long memory simply by virtue of its dynamics, as has been demonstrated in random (13, 15–17) and brain-like networks (18). This maintenance of information in system dynamics can act as a short-term memory and is a good candidate condition for consciousness. Indeed, previous studies have demonstrated altered chaoticity during tasks of relaxation and concentration (19), propofol-induced unconsciousness (2, 20), disorders of consciousness (2, 20) and in patients suffering from epilepsy (2, 21). In line with previous research (2, 20) we showed that chaoticity increased during propofol-induced unconsciousness. In addition, we were able to extend this finding to xenon-induced unconsciousness and demonstrated that the increase of chaoticity is specific to the loss of consciousness (i.e. using propofol and xenon), rather than the effect of altered neurochemistry per se (i.e. during exposure to ketamine).

*Edge of synchrony and consciousness*

A third critical phase transition that has been studied in the brain is the 'edge of synchrony', that is, the transition point between synchronous and asynchronous dynamics in a system of coupled oscillators. If phase synchrony between oscillating neural populations mediates information sharing (Varela, 2001; Fries, 2005), then the edge of synchrony provides a useful, dynamics-rich middle ground where transient coalitions of oscillators fall in and out of synchrony at every spatiotemporal scale—striking a balance of integration and segregation much in the same way as avalanche criticality (Yoon et al., 2015). Here, we analyzed the pair correlation function (PCF) in the alpha band, as well as narrowband detrended fluctuation analysis (see Supplementary Material), both of which have been discussed as measures of edge-of-synchrony criticality (12, 22). The PCF is a measure of the variability of oscillatory phase coupling, and has previously been used to demonstrate a link between criticality and integrated information (22) and its effect on pharmacologically-induced unconsciousness (23).

### Supplementary Methods, Note2

Several methods have been used to estimate the closeness of a given neuronal avalanche distribution to a power law (Shew et al., 2009; Klaus et al., 2011; Varley et al., 2020). Here, we propose a method based on the slope of the best-fit power law. For increasingly subcritical avalanche dynamics, the (absolute) slope will increase as large avalanches become increasingly rare. The magnitude of the increase in slope between two distributions thus quantifies the shift towards subcriticality. We thus fit the distribution to a power law using the Python 'powerlaw' package (Alstott et al., 2014). Specifically, we fit the probability distribution of avalanche size and duration to a truncated power law function with exponential cutoff, with an $x_{min}$ set to the minimum avalanche size and duration, respectively. The $x_{max}$ was optimized to yield the best fit. This is in contrast with the approach of Varley et al. (2020) who set both $x_{min}$ and $x_{max}$ to optimize best fit. Our rationale for fixing $x_{min}$ to the data minimum was to ensure that the lower (small x) part of the distribution was consistently used for slope estimation.

Note that the slope thus estimated is not in general equivalent to a critical exponent. Strictly speaking, critical exponents only characterize critical dynamics; away from criticality, this value still can serve as an indicator of the degree of subcriticality, but it loses its connection to the theory of critical exponents (see Sethna et al., 2001 for an excellent review).
